# A natural encoding of genetic variation in a Burrows-Wheeler Transform to enable mapping and genome inference

**DOI:** 10.1101/059170

**Authors:** Sorina Maciuca, Carlos del Ojo Elias, Gil McVean, Zamin Iqbal

## Abstract

We show how positional markers can be used to encode genetic variation within aBurrows-Wheeler Transform (BWT), and use this to construct a generalisation ofthe traditional “reference genome”, incorporating known variation within aspecies. Our goal is to support the inference of the closest mosaic of previously known sequences to the genome(s) under analysis.

Our scheme results in an increased alphabet size, and by using a wavelet tree encoding of the BWT we reduce the performance impact on rank operations. We give a specialised form of the backward search that allows variation-aware exact matching. We implement this, and demonstrate the cost of constructing an index of the whole human genome with 8 million genetic variants is 25GB of RAM. We also show that inferring a closer reference can close large kilobase-scale coverage gaps in *P. falciparum*.

## 1 Introduction

Genome sequencing involves breaking DNA into fragments, identifying substrings (called “reads”), and then inferring properties of the genome. Recently, it has become possible to study within-species genetic variation on a large scale [6, 7], where the dominant approach is to match substrings to the canonical “reference genome” which is constructed from an arbitrary individual. This problem (“mapping”) has been heavily studied (see [5]) and the Burrows-Wheeler Transform (BWT) [2] underlies the two dominant mappers [3, 4]. Mapping reads to a reference genome is a very effective way of detecting genetic variation caused by single character changes (SNPs - single nucleotide polymorphisms). However, this method becomes less effective the further the genome differs from the reference. This is an important problem to address since, in many organisms, biologically relevant genomic regions are highly diverse.

For a given species, our goal is to build a compact representation of the genomes of N individuals, which we call a Population Reference Genome (PRG). This data structure facilitates the following inference: we take as input sequence data from a new sample, an estimate of how many genomes the sample contains and their relative proportions - e.g. a normal human sample would contain 2 genomes in a 1:1 ratio, a bacterial isolate would contain 1 genome and a malaria sample might contain 3 genomes in the ratio 10:3:1. We would then infer the sequence of the underlying genomes. In this paper we describe a method for encoding genetic variation designed to enable this approach.

Genomes evolve mainly via two processes - mutation (changing a few characters) and recombination (either two chromosomes exchange a chunk of DNA, or one chromosome copies a chunk from another). Thus once we have seen many genomes of a given species, a new genome is likely to look like a mosaic of genomes we have seen before. If we can infer a close mosaic, we have found a “personalised reference genome”, and reads are more likely to match exactly. This approach was first described in [8], applied to the human MHC region. However their implementation was quite specific to the region and would not scale to the whole genome. Valenzuela *et al*. [1] have also espoused a find-the-closest-reference approach.

Other “reference graph” methods have been published [9, 10, 11], generally approaching just the alignment step. Siren *et al*. developed a method (GCSA [10]), with construction costs for a whole human genome (plus mutations) of more than 1 Tb of RAM. Huang *et al*. [11] developed an FM-index [13] encoding of a reference genome-plus-variation (“BWBBLE”) by extending the genetic alphabet to encode single-character variants with new characters and then concatenating padded indel variants to the end of the reference genome. We do something similar, but treat all variation in an equivalent manner, and retain knowledge of allelism naturally. While completing this paper, the preprint for GCSA2 was published ([12]), which drops RAM usage of human genome index construction to <100GB at the cost of>1Tb of disk I/O.

We show below how to encode a set of genomes, or a reference plus genetic variation, in an FM-index which naturally distinguishes alternate alleles. We extend the well known BWT backward search and show how read-mapping can be performed in a way that allows reads to cross multiple variants, allowing recombination to occur naturally. Our data structure supports bidirectional search (which underlies the Super Maximal Exact Match algorithms of bwa-mem [3]), but currently we have only implemented exact matching. We use empirical datasets to demonstrate low construction cost (human genome) and the value of inferring a personalised reference in *P. falciparum*.

## 2 Background: Compressed Text Indexes

### 2.0.1 Burrows-Wheeler Transform

The Burrows-Wheeler Transform (BWT) of a string is a reversible permutation of its characters. The BWT of a string *T* =*t*_1_ *t*_2_ … *t_n_* is constructed by sorting its *n* cyclic shifts *t*_1_ *t*_2_ … *t_n_*, *t*_2_ … *t_n_ t*_1_, …, *t_n_ t*_1_ … *t*_*n*−1_ in lexicographic order. The matrix obtained is called the Burrows-Wheeler Matrix (BWM) and the sequence from its last column is the BWT.

### 2.0.2 Suffix Arrays

The suffix array of a string *T* is an array of integers that provides the starting position of *T*’s suffixes, after they have been ordered lexicographically. Formally, if *T_i,j_* is the substring *t_i_t*_*i*+1_ …*t_j_* of *T* and SA is the suffix array of *T*, then *T_SA[1],n_* < *T_SA[2],n_* < … < *T_SA[n],n_*. It is related to the BWT, since looking at the substrings preceding the terminating character $ in the BWM rows gives the suffixes of *T* in lexicographical order.

### 2.0.3 Backward search

Any occurrence of a pattern *P* in text is a prefix for some suffix of *T*, so all occurrences will be adjacent in the suffix array of *T* since suffixes starting with *P* are sorted together in a SA-interval. Let *C* [*α*] be the total number of occurrences in *T* of characters smaller than *a* in the alphabet. If *P′* is a suffix of the query *P* and [*l*(*P′*), *r*(*P′*)) is its corresponding SA-interval, then the search can be extended to *aP′* by calculating the new SA-interval:

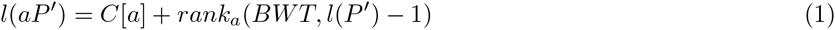

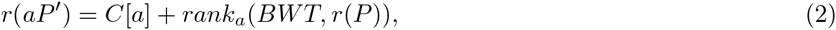

where the operation *rank_a_*(*S*, *i*) returns the number of occurrences of symbol *a* in *S*[1, *i*]. The search starts with the SA-interval of the empty string, [1, *n*] and successively adds one character of *P* in reverse order. When the search is completed, it returns a SA-interval [*l*, *r*) for the entire query *P*. If *r* > *l*, there are *r* − *I* matches for *P* and their locations in *T* are given by SA[*i*] for *I* ≥ *i* < *r*. Otherwise, the pattern does not exist in *T*. If the *C*-array and the ranks have already been stored, the backward search can be performed in *O*(|*P*|) time in strings with DNA alphabet.

### 2.0.4 Wavelet Trees

Rank queries scale linearly with the alphabet size by default. The wavelet tree [14] is a data structure designed to store strings with large alphabets efficiently and provide rank calculations in logarithmic time. The tree is defined recursively: take the lexicographically ordered alphabet, split it into 2 equal halves; in the string corresponding to the current node (start with the original string at root), replace the first half of letters with 0 and the other half with 1; the left child node will contain the 0-encoded symbols and the right child node will contain the 1-encoded symbols, preserving their order from the original string; re-apply the first step for each child node recursively until the alphabet left in each node contains only one or two symbols (so a 0 or 1 determines which symbol it is).

To answer a rank query over the original string with large alphabet, repeated rank queries over the bit vectors in the wavelet tree nodes are used to locate the subtree that contains the leaf where the queried symbol is non-ambiguously encoded. The rank of the queried symbol in this leaf is equal to its rank in the original string. The number of rank queries needed to reach the leaf is equal to the height of the tree, i.e. log_2_ |Σ| if we let Σ be the set of symbols in the alphabet. Computing ranks over binary vectors can be done in constant time, so a rank query in a wavelet tree-encoded string has complexity *O*(log_2_ |Σ|).

## 3 Encoding a variation-aware reference structure

### 3.1 Terminology

A *variant site* or *site* is a region of the chromosome where there are a number of alternative options for what sequence can be present. These alternatives are termed *alleles* and might be as short as a single character, or could be many hundreds of characters long. A *pan-genome* refers to a representation (with unspecified properties) of a number (greater than 1) of genomes within a species. A Population Reference Graph is an encoding of a pan-genome that enables matching of sequence data to the datastore, inference of nearest mosaic with the appropriate ploidy, and then discovery of new variants not present in the PRG.

**Figure 1.**
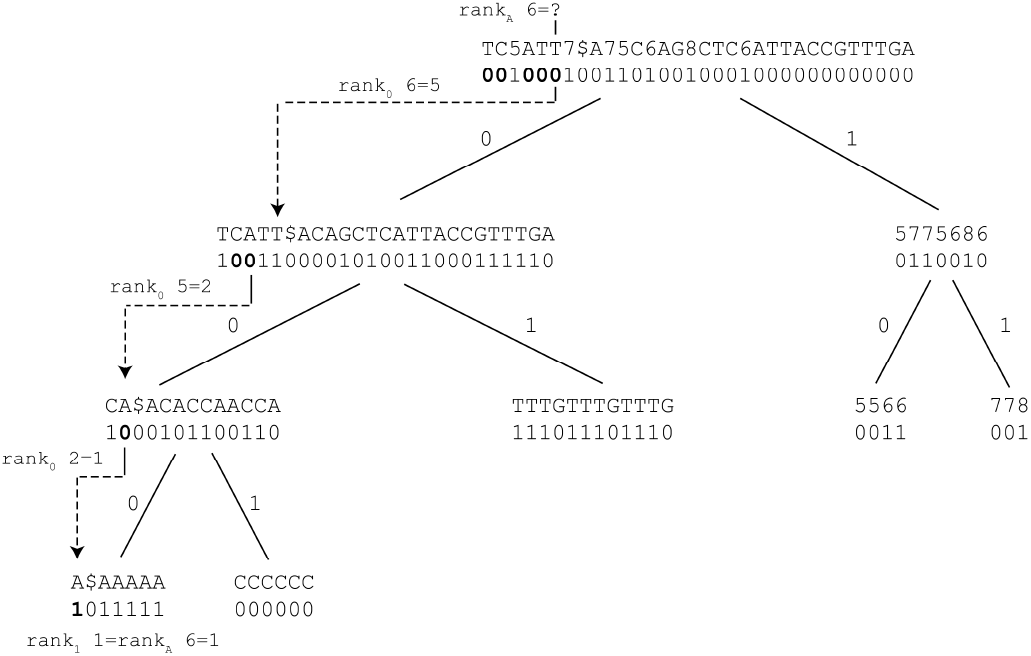
Wavelet tree encoding of a string that is the same as the BWT in figure 4. Calculating the rank of the marked “A” is performed by repeated rank() calls moving down the binary tree until the alphabet remaining is just 2 characters. Note that only the bit vectors are stored in the tree, the corresponding strings are only shown here for clarity.

**Figure 2.**
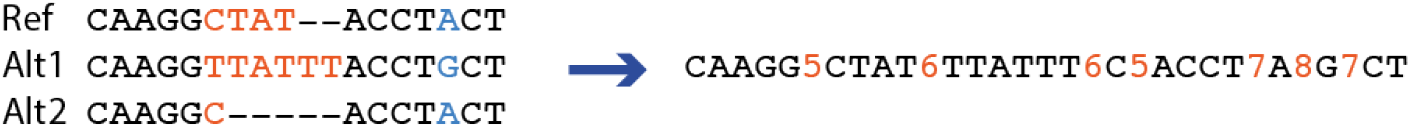
A simple PRG linearised according to our encoding. The first site has 3 alleles, which do not here look at all similar, and the second is a SNP.

### 3.2 PRG Encoding

We use a PRG conceptually equivalent to a directed, acyclic, partial order graph, that is generated from a reference sequence and a set of alternative sequences at given variation sites. The graph is linearised into a long string over an alphabet extended with new symbols marking the variants, for which the FM-index can be constructed. We call this string the *linear PRG*.

Building this data structure requires multiple steps.

1. Corresponding regions of shared sequence between the input genomes must be identified. These must be of size *k* at least (where *k* is pre-defined), and act as anchors.
2. For any site between two anchor regions, the set of possible alleles/haplotypes must be determined, but do not need to be aligned. Indels are supported by haplotypes of different lengths.
3. Each variation site is assigned two unique numeric identifiers, one even and one odd, which we call variation markers. The odd identifiers will mark variation site boundaries and will sometimes be referred to as site markers. The even identifiers will mark alternative allele boundaries and will sometimes be referred to as allele boundary markers.
4. For each variation site, its left anchor is added to the linear PRG, followed by its odd identifier. Then each sequence coming from that site, starting with the reference sequence, is successively added to the linear PRG, followed by the even site identifier, except the last sequence, which is followed by the odd identifier.
5. Convert the linear PRG to integer alphabet (*A* → 1, *C* → 2, *G* → 3, *T* → 4, variation site identifiers → 5,6,…)
6. The FM-index (suffix array, BWT, wavelet tree over BWT) of the linear PRG is constructed and we will call this the vBWT.

**Figure 3.**
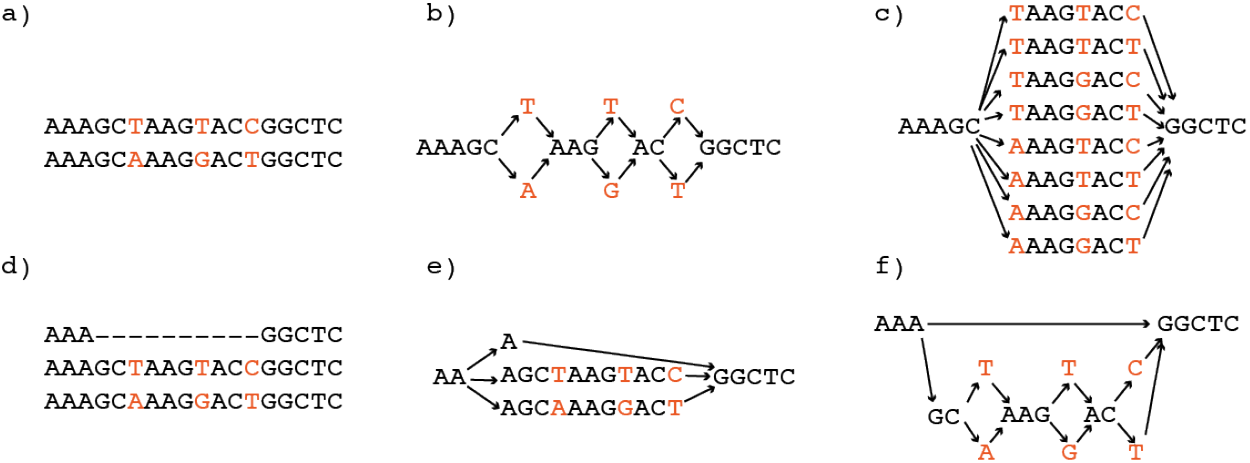
PRG graph structure. The sequences shown in Figure 3a) could be represented either as 3 separate mutations (shown in b)), or enumerated as 8 small haplotypes, shown in c). Both are supported by our encoding. Similarly, the sequences in d) could be represented in our implementation as shown in e). However, we do not support “nesting” of alleles, as shown in f).

An illustration of these steps on a toy example is given in Figure 2.

Importantly, **the markers force the ends of alternative sequences coming from the same site to be sorted together in a separate block in the Burrows-Wheeler matrix, even if they do not have high sequence similarity**. Therefore, alternative alleles from each site can be queried concurrently.

### 3.3 Graph structure: constraints

We show in Figure 3a) two sequences which differ by 3 SNPs and give two graph encodings in 3b) and 3c). Both represent the sequence content equally well, and we allow both. In 3d) we have an example where a long deletion lies “over” two other alleles. We would encode this in our PRG as shown in 3e). This works but results in many alternate alleles. An alternative would be to allow “nested” variation, where variants lie on top of other alleles, as shown in Figure 3f). This could be encoded in our system, but we do not allow it for our initial implementation, as it would potentially impact mapping speed.

## 4 Variation-aware backward search in vBWT

In this section, we present a modified backward search algorithm for exact matching against the vBWT that is aware of alternative sequence paths. Our implementation leverages the succinct data structures library SDSL [18] and is incorporated in our software called **gramtools**.

**Figure 4.**
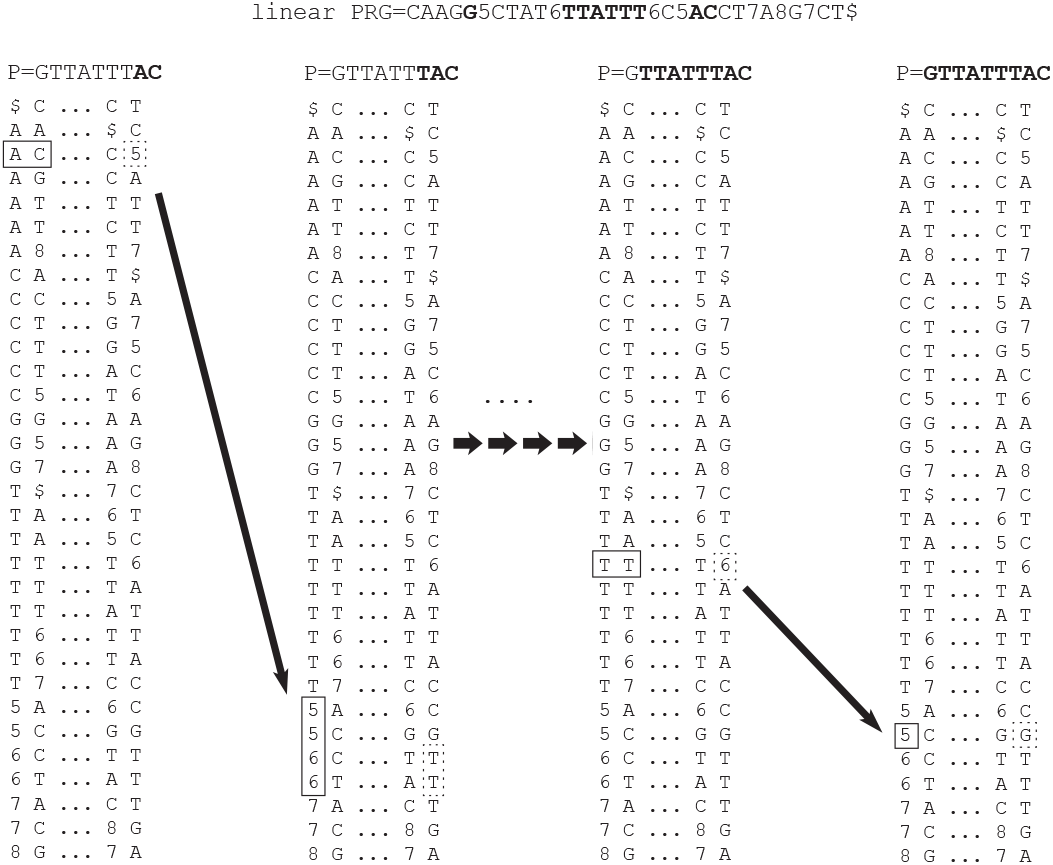
Backward search across the vBWT of the linear PRG in figure 2. We start at the right-hand end of the read GTTATTTAC, with the character C, and as we extend we hit the character 5, signalling the start or end of a variation site. We check the suffix array to get the coordinate in the linear PRG, and find it is the end. Therefore, the read must now continue into one of the alleles, signalled by the number 6. Continuing in this manner (the shorter arrows signify multiple intermediate steps not shown) we are able to align across the site.

When reads align to the non-variable part of the PRG or when an allele is long enough to enclose the entire read, the usual backward search algorithm can be used. Otherwise, when the read must cross variation site junctions, site identifiers and some alternative alleles must be ignored by the search. This means a read can align to multiple substrings of the linear PRG that may not be adjacent in the BWM, so the search can return multiple SA-intervals. We give pseudocode in Algorithm 1 below, and outline the idea in Figure 4.

At each step in the backward search, before extending to the next character, we need to check whether the current matched read substring is preceded by a variation marker anywhere in the linear PRG. A scan for symbols larger than 4 (“range_search_2d” in the pseudocode) must be performed within the range given by the current SA-interval. With a wavelet tree this range search can be done in *O*(*d*log(|Σ|/*d*)) time, where d is the size of the output. If a variation marker is found and it is an odd number, the read is about to cross a site boundary. The suffix array can be queried to find the position of the two odd numbers (start/end of the site) in the linear PRG.

If the search cursor is next to the start of the site, it is just the site marker that needs to be skipped, so the SA-interval (size 1) of the suffix starting with that marker needs to be added to the set of intervals that will be extended with the next character in the read. If the search cursor is next to the end of a site, all alternative alleles from that site need to be queried. Their ends are sorted together in the BWM because of the markers, so they can be queried concurrently by adding the SA-interval of suffixes starting with all numbers marking that site (even and odd).

### Algorithm 1 Variation-aware backward search

~~~
          **Input:** pattern P[1, m] and FM-index of PRG in integer alphabet
            **Output:** list of SA intervals corresponding to matches of P
1: *l* → *C*(*P*[*m*])                 ⊳l for left
2: *r* → *C*(*P*[*m*] + 1)                                       ⊳ r for right
3: *i* → *m*
4: SA_int ={[*l*, *r*)}                                   ⊳ list of SA intervals
5: Extra_int = 0              ⊳ Extra intervals
6: **while** *i* > 1 and SA_int = 0 **do**
7:    **for all** [*l*, *r*) ∈ SA_int **do**
8:     *M* → WT.range_search_2d(*l*, *r* − 1, 5, |Σ|)       ⊳ find variation site markers
9:     *for all* (*idx*, *num*) ∈ *M* **do**             ⊳*idx € [l,r),num*€ [5, |Σ|]
10:          **if** *num*%2 = 0 **then**
11:                              odd_num = num − 1
12:          **else**
13:        odd_num = *num*
14:         **if** *SA*[*C*(odd_num)] < *SA*[*C*(odd_num) + 1] **then**
15:          start_site → *C*(odd_num), end_site → *C*(odd_num) + 1
16:         **else**
17:          start_site → *C*(odd_num) + 1, end_site → *C*(odd_num)
18:        **if** *num*%2 = 1 **and** *SA*[*idx*] = *SA*[end_site] + 1 **then**
19:         Extra_int = Extra_int ∪ {[*C*(*num*), *C*(*num* + 2)]}
20:         **else**
21:              Extra_int = Extra_int ∪ {[*C*[start_site], *C*[start_site] + 1]}
22:  *i* → *i* − 1
23: SA_int = SA_int ∪ Extra_int
24:    **for all** [*l*, *r*) ∈ SA_int **do**
25:    *l* = *C*(*P* [*i*]) + *rank_BWT_* (*P* [*i*],*l* − 1)
26:    *r* = *C*(*P* [*i*]) + *rank_BWT_* (*P* [*i*],*r*)
~~~

If the variation marker found is even, the read is about to cross an allele boundary, which means its current suffix matches the beginning of an alternative allele and the read is about to walk out of a site, so the search cursor needs to jump to the start of site. The odd markers corresponding to that site can be found in the first column of the BWM, and then querying the suffix array decides which one marks the start of site. The SA-interval (size 1) for the BWM row starting with this odd marker is recorded. Once the check for variation markers is finished and all candidate SA-intervals have been added, each interval can be extended with the next character in the read by using equations 1 and 2.

## 5 Experiments

### 5.1 Construction cost : the human genome

We constructed a PRG from the human reference genome (GRC37 without “alt” contigs) plus the 1000 genomes final VCF (12GB in size) [6]. We excluded variants without specified alleles, and those with allele frequency below 5% (rare variation offers limited benefit - our goal is to maximise the proportion of reads mismatching the graph by at most 1 SNP). If two variants occurred at consecutive bases, they were merged, and all haplotypes enumerated. If the VCF contained two consecutive records which overlapped, the second was discarded. This resulted in a dataset of 7.4 million SNPs and 978000 indels. We give construction costs in Table 1, along with comparative figures for BWBBLE with identical input.

For comparison, GCSA took over 1TB of RAM building chromosomes separately and pruning the graph in high diversity regions. GCSA2 reduces the memory footprint to below 128GB RAM, running in 13 hours with 32 cores, and using over 1Tb of I/O to fast disk. Our vBWT construction has a lower memory cost than GCSA, GCSA2 and BWBBLE, is faster than GCSA/GCSA2, has no (significant) I/O burden, but is significantly slower than BWBBLE.

**Table 1.** FM-index construction costs and final data structure size for human reference genome plus 1000 genomes variants

| Software | Peak Memory(GB) | Time (hrs/mins) | Final/in-use memory (GB) |
| --- | --- | --- | --- |
| vBWT | 25 | 4h24m | 17.5 |
| BWBBLE | 60 | 1h5m | 12 |

**Figure 5.**
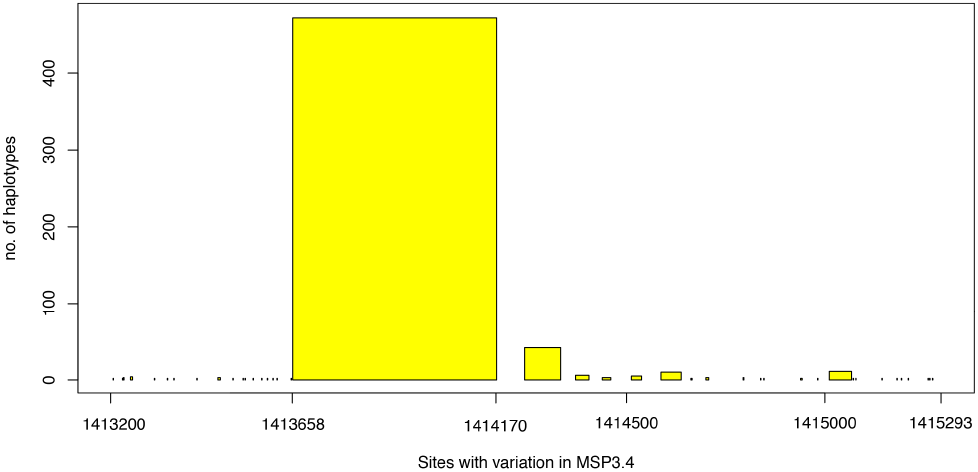
Histogram of number of alleles at each site in MSP3.4 plotted above the chromosome coordinate.)

### 5.2 Inferring a Closer Reference Genome

*P. falciparum* is a haploid parasite that undergoes recombination. It has an unusual genome that contains more indels than SNPs [15]. The gene MSP3.4 is known to have two diverged lineages at high frequencies in multiple populations from across the world. The lineages differ by around 1 SNP every 3 bases over a 500bp region (the DBL domain) of the gene. We constructed a catalog of MSP3.4 variation from Cortex [16] variant calls from 650 *P. falciparum* samples and built a PRG just for that chromosome. We show in Figure 5 the density of variants and number of alleles.

We aligned Illumina 76bp reads from a well-studied sample that was not used in graph construction (named 7G8) to the PRG using backward search (exact matching, which took 3 mins), and collected counts on the number of reads supporting each allele. At each site we chose the allele with the highest coverage to extract the path through the graph with maximum support - this was our graph-inferred personalised reference for this sample. We then mapped the reads (using bwa_mem [17]) independently to the reference and to the inferred genome. As can be seen in Figures 6 and 7, our method gives dramatically better pileup results over the MSP3.4 gene.

### 5.3 Simulations, usability, future performance improvements

We took 44,439 *P. falciparum* SNPs and indels called with Cortex from a single genetic cross (7G8xGB4) [15] and created a whole-genome PRG, and simulated 10,000 reads from one random haplotype. All reads were 150bp, error-free. We precalculate a hash of the SA intervals corresponding to all 9-mers in the PRG that overlap a variation site. This one-time precalculation was done in 1 hour 43 mins using 25 threads. In order to avoid unfairly slowing BWBBLE we constrained it to do exact matching only. All experiments were performed single-threaded on a machine with 64 processors Intel Xeon CPU E5-4620 v2 @ 2.60GHz and 1 TB of memory.

**Figure 6.**
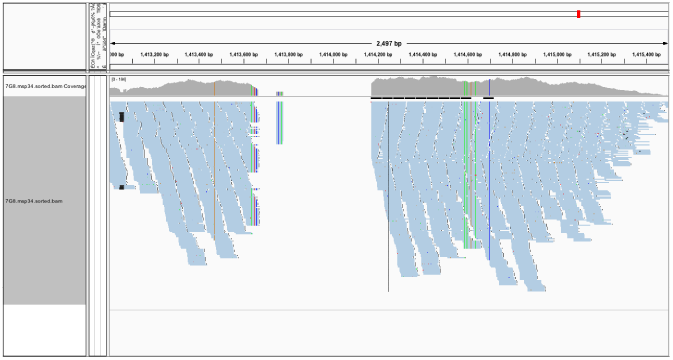
Mapping reads from sample 7G8 to *P. falciparum* 3D7 reference genome results in a gap covering the DBL domain

**Figure 7.**
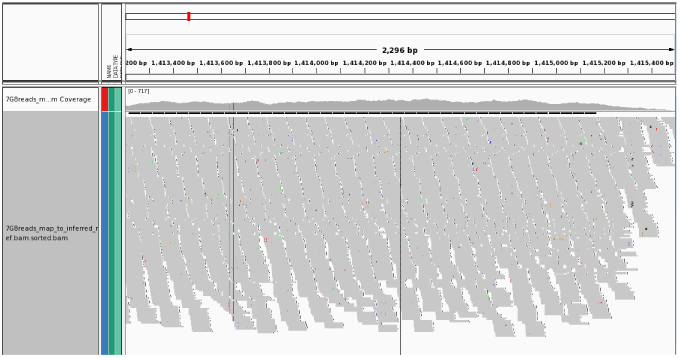
Mapping reads from sample 7G8 to our vBWT-inferred genome removes the gap, leaving isolated variants easy to detect with standard methods

**Table 2.** Simulation results

| Software | Step | Time (sec) | Speed (reads/sec) |
| --- | --- | --- | --- |
| gramtools | Map and Infer reference | 1051 | 9.5 |
| gramtools | bwa map to ref | 0.861 | 11614 |
| BWBBLE | align | 2.6 | 3846 |

Gramtools mapping speed is notably slower than BWBBLE, although it is usable for megabase sized genomes - a 30x whole-genome dataset for *P. falciparum* would take 5.8 hours using 24 cores. However, the output is directly usable and interpretable by any bioinformatician - a reference genome close to the sample, and a standard SAM file. By comparison, BWBBLE outputs a SAM file with respect to an artificial reference with indels appended at the end - to use this in a normal pipeline requires software development and innovation.

There are a number of performance improvements we can make. We store an integer array that allows us to determine if a position in the PRG is in a site, and if so, which allele; this is naively encoded (in std::vector). For the human example, this costs us around 12GB of RAM. This array, which contains a zero at every non-variable site in the chromosome, could be stored much more compactly. More significantly, there is one significant speed improvement which we have yet to implement - precalculating and storing an array of ranks at marker positions across the BWT - just as in a standard FM-index. This is not normally done for large alphabets, but we can store only for A,C,G,T.

## 6 Discussion

We have described a whole-genome scale implementation of a PRG designed to enable inference of a within-graph mosaic reference close to that of a new individual, followed by discovery of novel variation as a “delta” from that. As with any reference graph approach, there is an implicit coupling between mapping and graph structure (for handling alternate alleles). By placing positional markers, we are able to ensure that alternate alleles sort together in the BWT matrix, allowing mapping across sites and recombination. For haploids we naturally infer a personalised reference genome. For other ploidies, our implementation readily lends itself to “lightweight alignment” [19, 20, 21] followed by an HMM to infer haplotypes, followed by full MEM-based graph alignment.

### 6.0.1 Software

Our software, gramtools, is available here: http://github.com/iqbal-lab/gramtools.

## Acknowledgements

We would like to thank Jacob Almagro-Garcia, Phelim Bradley, Rayan Chikhi, Simon Gog, Lin Huang, Jerome Kelleher, Heng Li, Gerton Lunter, Rachel Norris, Victoria Popic, and Jouni Siren for discussions and help. We thank the SDSL developers for providing a valuable resource.

